# An integrative machine learning approach identifies the centrality of ferroptosis, cuproptosis, and immune pathway crosstalk for breast cancer stratification and therapy guidance

**DOI:** 10.1101/2025.09.08.674809

**Authors:** Syed Ahsan Shahid, Ahmed Al-Harrasi, Adil Alsiyabi

## Abstract

Breast cancer (BRCA) is a leading cause of cancer-related mortality in women, characterized by marked heterogeneity in molecular subtypes, immune microenvironment, and therapeutic response. Current gene expression classifiers often lack mechanistic grounding, limiting their clinical utility. Using an integrated machine learning approach, we identified a four-gene panel, *FOXO4, EGFR, FGF2,* and *CDKN2A*, capturing convergent dysregulation across ferroptosis, cuproptosis, and immune pathways. The panel reflects not only redox and proliferative dysregulation but also distinct immune microenvironmental patterns, with *FGF2* linked to stromal remodeling and *CDKN2A* correlated with adaptive immune responses, underscoring its biological integration into BRCA pathology. This panel is rooted in recurrent dysregulation of oxidative stress control *(FOXO4)*, proliferative and angiogenic signalling *(EGFR, FGF2)*, and cell-cycle-immune interfaces *(CDKN2A)*, linking classification to central regulatory mechanisms. The model achieved 97-98% test accuracy (AUC 0.99) for tumour-healthy discrimination. Our findings reveal transcriptional convergence between redox-metabolic and immune-escape programs in BRCA. We propose a compact and interpretable panel with translational potential, offering a minimal, mechanism-informed diagnostic framework for clinical deployment in breast cancer for precision oncology.

## 1 Introduction

Breast cancer (BRCA) remains the most frequently diagnosed malignancy and the leading cause of cancer-related mortality among women globally, with over 2.3 million new cases and approximately 685,000 deaths reported in 2020 [1]. Despite significant advances in molecular diagnostics and targeted therapeutics, BRCA continues to pose a major clinical challenge due to its marked heterogeneity, therapeutic resistance, and variation in immune and metabolic profiles across molecular subtypes [2]. Gene expression-based classifiers have supported risk stratification and treatment selection but often suffer from redundancy, limited mechanistic relevance, and reduced generalizability across patient cohorts [3]. Therefore, there is a pressing need to develop compact, biologically informed gene signatures that enable accurate tumor classification while capturing critical aspects of tumor biology and therapeutic vulnerability.

Evidence increasingly supports a key role for regulated cell death pathways, such as ferroptosis and cuproptosis, alongside immune modulation, in shaping tumor progression and response to therapy [4–7]. Ferroptosis, a non-apoptotic iron-dependent form of cell death driven by lipid peroxidation and oxidative imbalance, has shown tumor-suppressive effects in aggressive BRCA subtypes [8, 9]. Cuproptosis, a recently defined copper-dependent mechanism linked to mitochondrial metabolism and protein lipoylation, represents a distinct vulnerability in tumors with disrupted energy metabolism [10, 11]. Additionally, the tumor immune microenvironment, influenced by cytokine signaling, immune checkpoints, and antigen presentation, plays a pivotal role in tumor progression and therapeutic response [12–14]. Each of these regulatory axes, ferroptosis, cuproptosis, and immune activity, has been individually implicated in BRCA. However, no prior study has integrated them into a unified minimal gene panel for both classification and biological insight. Publicly available transcriptomic datasets, such as those from The Cancer Genome Atlas (TCGA) [15], offer opportunities to identify such convergences through computational approaches. However, many existing classifiers prioritize statistical performance over biological interpretability, often relying on large gene sets with limited mechanistic anchoring. Moreover, high-dimensional models also present barriers to clinical implementation, particularly in resource-limited settings or streamlined diagnostic workflows.

To address this gap, we developed a four-gene classifier for BRCA based on the convergence of ferroptosis, cuproptosis, and immune signaling. Using a multi-model machine learning framework incorporating LASSO, Random Forest, XGBoost, and SHAP interpretability, we refined a nine-gene input list of differentially expressed genes, curated from validated regulators of ferroptosis, cuproptosis, and immune pathways, into a compact four-gene panel. Tumor immune contexture was assessed using xCell-based deconvolution across 64 immune and stromal cell types in over 1,200 BRCA samples. The developed workflow resulted in a high-performance diagnostic panel mechanistically embedded in core oncogenic and immunological pathways. Each selected gene reflects distinct functional processes spanning oxidative stress, proliferative signaling, and immune interaction, enabling classification informed by underlying regulatory architecture rather than surface expression patterns alone. Such interpretable, compact signatures are particularly valuable where broad transcriptomic profiling is impractical. Thus, we propose a biologically interpretable, clinically feasible four-gene BRCA classifier derived from the intersection of ferroptosis, cuproptosis, and immune regulation. This integrative approach advances molecular classification by prioritizing biological context as the driver of dimensionality reduction and biomarker discovery.

**Figure 1:**
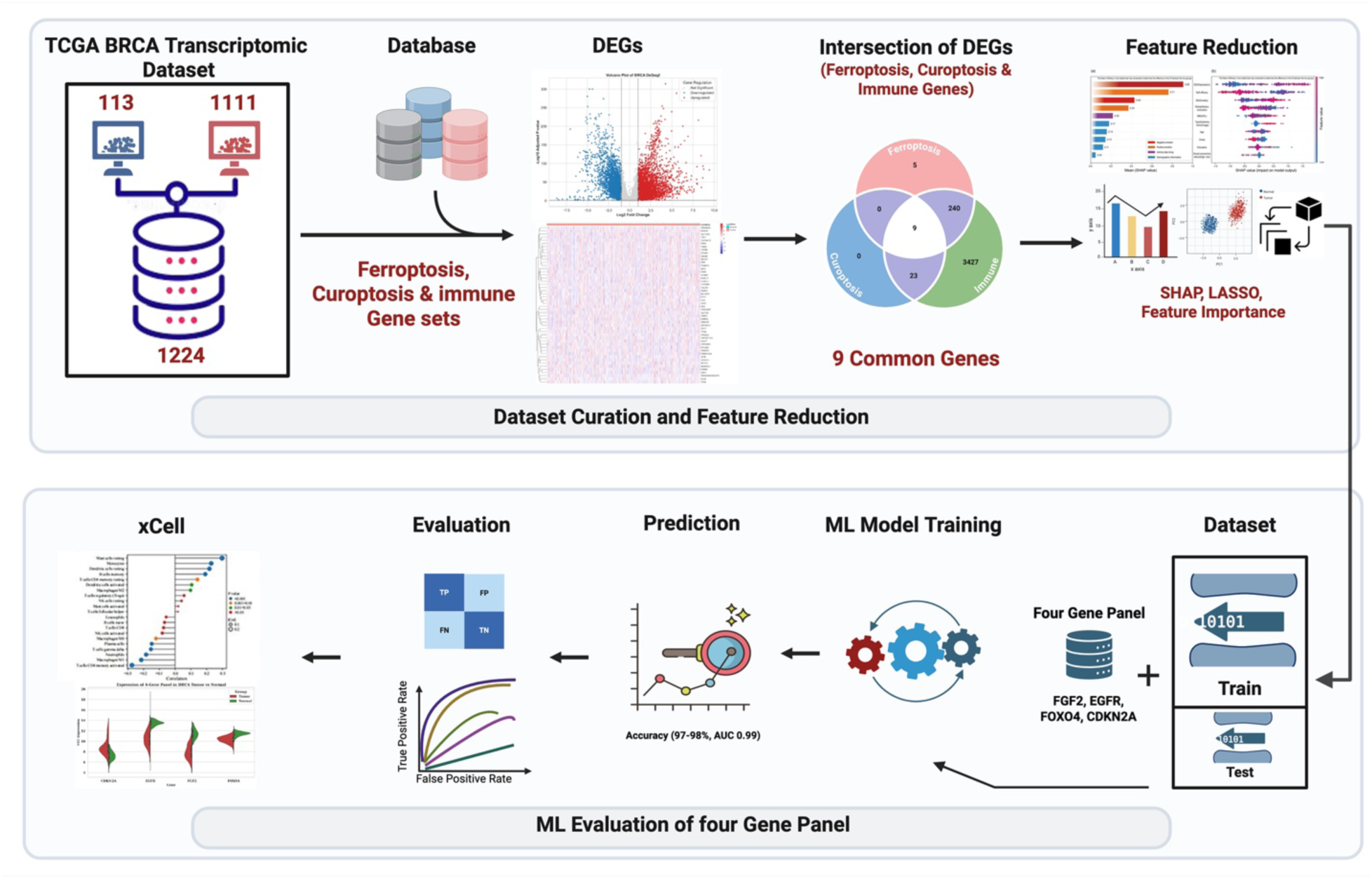
Workflow for identification and evaluation of a four-gene BRCA classifier panel.

## 2 Methodology

### 2.1 Transcriptomic data acquisition and preprocessing

RNA-sequencing data for breast invasive carcinoma (BRCA) were retrieved from The Cancer Genome Atlas (TCGA) [15] via the Genomic Data Commons (GDC) portal using *TCGAbiolinks*. Gene-level raw counts from the STAR alignment pipeline (“Gene Expression Quantification” workflow) were obtained, comprising 1,111 primary tumors and 113 solid tissue normals (n = 1,224). To minimize memory load and download errors, samples were retrieved in barcode-based batches of ∼100 and parsed into *SummarizedExperiment* objects. Fault-tolerant downloading was applied, ensuring incomplete or corrupted chunks were skipped. Metadata were harmonized, and final *SummarizedExperiment* objects were merged into a unified dataset containing expression counts and annotations. The consolidated matrix and metadata were archived in standard formats for downstream analyses.

### 2.2 Differential gene expression analysis

Differential gene expression analysis was performed using the DESeq2 package (v1.40.2) [16]. Raw gene-level count data and matched sample annotations were curated to ensure accurate alignment between tumor and normal samples, based on TCGA BRCA metadata. Genes with fewer than 10 counts in at least 10% of samples were excluded to reduce background noise and improve statistical power. The filtered count matrix was encapsulated into a *DESeqDataSet* object, with tumor versus normal tissue specified as the primary contrast. The standard DESeq2 pipeline was executed, and log₂ fold change estimates were refined using the *apeglm* shrinkage method to improve interpretability of effect sizes. Ensembl gene identifiers were converted to HGNC symbols using biomaRt (v2.56.1) [17], and unmappable entries were filtered out. Genes were classified as differentially expressed if they satisfied both an adjusted *P*-value threshold of < 0.05 and an absolute log2 fold change > 1. The top 50 most significantly differentially expressed genes (DEGs), ranked by adjusted *P*-value, were visualized using clustered heatmaps of svariance-stabilized expression values. To provide a transcriptome-wide overview of changes, a volcano plot was generated, highlighting significantly up- and downregulated genes. Complete DEG results and filtered outputs were archived for downstream functional enrichment and modeling analyses.

### 2.3 Variance-stabilizing transformation and gene symbol annotation

To correct for differences in sequencing depth and reduce heteroscedasticity in expression measurements, a variance-stabilizing transformation (VST) was applied to the raw count data using the DESeq2 package. Both tumor and normal samples, as defined by TCGA BRCA metadata, were incorporated into the normalization workflow to preserve condition-specific variance. Library size differences were corrected by estimating size factors prior to transformation. VST was performed with a *blind* parameter set to *FALSE* to retain the biological signal relevant to group comparisons. Gene annotation was carried out by mapping Ensembl gene identifiers to HGNC gene symbols via the Ensembl BioMart interface. Ensembl version suffixes were stripped prior to mapping. Genes without valid HGNC annotations were excluded. For cases in which multiple Ensembl IDs mapped to a single gene symbol, expression values were averaged to yield a unique value per gene. The resulting matrix, indexed by HGNC symbols, represented a robust, normalized expression dataset suitable for downstream analyses, including gene selection, functional enrichment, and classification modeling.

### 2.4 Curation of ferroptosis-, cuproptosis-, and immune-related gene sets

Gene sets relevant to ferroptosis, cuproptosis, and immune regulation were systematically curated to inform downstream transcriptomic filtering and functional analysis. Ferroptosis-associated genes were obtained from FerrDb v2 (http://www.zhounan.org/ferrdb), a manually curated database cataloguing regulators of ferroptosis classified as drivers, suppressors, markers, or unclassified [18]. Only genes with valid HGNC symbols were retained, and those appearing in multiple categories were preserved with all associated annotations. This yielded a non-redundant list of 1,293 unique ferroptosis-related genes. Cuproptosis-related genes were derived from a recent pan-cancer network-based study by Pham et al. (2024), which systematically identified 133 genes associated with cuproplasia across 23 tumor types, including breast cancer. The gene set encompasses core regulators of copper-induced cell death as well as functionally co-expressed partners involved in copper transport, redox regulation, and mitochondrial metabolism [19]. Genes were curated using HGNC-standard nomenclature and retained in full without further filtering. For immune-related signatures, the C7: Immunologic Signatures gene set collection was downloaded from the Molecular Signatures Database (MSigDB v2025.1) (https://www.gsea-msigdb.org/gsea/msigdb) [20]. The full C7 matrix file (GMT format) was parsed to extract unique HGNC symbols representing a wide range of immune cell types, activation states, and cytokine-mediated responses. Together, these ferroptosis-, cuproptosis-, and immune-related gene sets provided a targeted biological framework for expression profiling, differential analysis, and pathway enrichment in the context of breast cancer transcriptomics.

### 2.5 Wilcoxon rank-sum test for ferroptosis-, cuproptosis-, and immune-related gene expression

To assess differential expression between tumor and normal tissues, we applied the non-parametric Wilcoxon rank-sum test to variance-stabilized expression values derived from DESeq2 (v1.40.2). Tumor and normal sample labels were defined based on TCGA BRCA metadata. Analyses focused on the curated ferroptosis, cuproptosis, and immune-related gene sets described in Section 2.4 [19]. Each gene within these sets was independently tested for expression differences between tumor and normal samples. Median expression levels were calculated for each condition, and log2 fold changes were estimated as the difference in group medians. P-values were adjusted using the Benjamini-Hochberg procedure to control the false discovery rate. Genes were considered significantly dysregulated if they exhibited an adjusted P-value < 0.05 and an absolute log2 fold change > 1. For each category, ferroptosis, cuproptosis, and immune-related, the top 30 most significant genes (ranked by FDR) were visualized using row-scaled heatmaps of VST-normalized expression values. Gene-level heatmaps were generated using the pheatmap package (v1.0.12) [21], with samples annotated by tumor/normal classification and clustering applied across genes. All statistical procedures were conducted in R (v4.3.1), and all intermediate and final outputs, including Wilcoxon statistics, significant gene lists, and heatmaps, were archived for reproducibility and downstream interpretation.

### 2.6 Identification of intersecting dysregulated genes across ferroptosis, cuproptosis, and immune programs

To explore potential regulatory convergence, we identified genes commonly dysregulated across the ferroptosis-, cuproptosis-, and immune-related databases defined in Section 2.4. For each gene set, significantly altered genes were selected based on Wilcoxon test results, using an adjusted *P*-value threshold of < 0.05 and an absolute log2 fold change > 1 (see section 2.5). Overlaps among the three gene lists were computed in R using set-based operations. Venn diagrams were generated to visualize intersections and facilitate interpretation. Genes shared across all three categories were considered candidate regulatory hubs and were prioritized for downstream analysis.

### 2.7 Machine learning-based gene prioritization and dimensionality reduction

To reduce the nine-gene intersection signature to a more compact and diagnostically efficient subset, we employed a machine learning–driven feature prioritization framework. Expression values for the nine intersecting genes were extracted from the VST-normalized matrix and standardized via z-score transformation. Binary labels (“Primary Tumor” = 1, “Solid Tissue Normal” = 0) were assigned based on TCGA metadata. Three interpretable classifiers, LASSO logistic regression, Random Forest, and XGBoost, were trained on the full nine-gene panel. Feature importance was assessed using model-specific criteria: absolute L1-penalized coefficients (LASSO) [22], Gini importance (Random Forest) [23], and gain score (XGBoost) [24]. To validate model-agnostic feature contributions, SHapley Additive exPlanations (SHAP) values were also computed for the XGBoost model [25]. A bootstrapped approach was used to quantify predictive performance as a function of gene set size. For each model, genes were ranked by importance, and iterative subsets ranging from the top 1 to all 9 genes were used to train classifiers across 100 stratified resampling iterations (80% training / 20% testing). Mean accuracy at each subset size was tracked to determine the minimal number of genes required to maintain robust classification performance. This approach enabled biologically grounded, data-driven dimensionality reduction, identifying the smallest high-performing gene subset for downstream use in classification modeling, interpretation, and panel design.

### 2.8 Classification of BRCA samples using a minimal gene panel

To assess the predictive performance of the minimal gene panel (FOXO4, EGFR, FGF2, CDKN2A), we implemented a machine learning classification pipeline across multiple commonly used algorithms. Expression data were obtained from the BRCA VST-normalized matrix and filtered to retain only the selected genes. Sample labels were curated from associated clinical metadata, with "Primary Tumor" and "Solid Tissue Normal" annotated as "Tumor" and "Normal" classes, respectively. All gene symbols were standardized to uppercase to ensure consistency in downstream analyses. The expression matrix was transposed, and the input features were standardized using *StandardScaler*. Class labels were binarized using *LabelEncoder*, encoding “Normal” as 0 and “Tumor” as 1. To address class imbalance, synthetic oversampling was applied using the SMOTE algorithm after splitting the dataset and before model training. Eleven classification models were benchmarked: Random Forest (RF), Support Vector Machine (SVM), XGBoost (XGB), AdaBoost (ADA), Gradient Boosting (GBM), k-Nearest Neighbors (KNN), Logistic Regression (LOGIT), Decision Tree (TREE), Gaussian Naive Bayes (NB), Multilayer Perceptron (ANN), and L1-penalized Logistic Regression (LASSO). Each model was trained and evaluated over 100 bootstrapped iterations using *StratifiedShuffleSplit* with an 80:20 train-test ratio, maintaining label distribution across splits. For each split, the pipeline (SMOTE + classifier) was fitted on training data and evaluated on both the train and test sets. Performance metrics, including accuracy, F1-score, precision, recall, ROC AUC, sensitivity, and specificity, were computed and recorded for each bootstrap iteration. Final model performance was summarized by computing the mean and standard deviation of each metric across all bootstraps. Results were exported as an aggregated performance matrix in Excel format for further comparative evaluation.

### 2.9 PCA and Gene Expression Visualization of Minimal Gene Panel

To further assess the biological relevance and discriminatory capacity of the 4-gene panel, we performed both violin plot-based expression comparison and principal component analysis (PCA) on BRCA samples. Variance-stabilized expression values were used for both analyses. For expression comparison, gene-wise violin plots were generated to visualize the distribution of each gene’s expression across tumor and normal samples, highlighting potential differential expression patterns. For dimensionality reduction, PCA was performed using the scikit-learn package (v1.3.0). Expression values for the four selected genes were standardized and projected onto the first two principal components. Samples were colored by group (Tumor vs Normal) to visualize class separability. The proportion of variance explained by each principal component was indicated on the axes. All plots were generated using Seaborn (v0.12.2) [26] and Matplotlib (v3.7.2) [27].

### 2.10 Immune Cell Infiltration Analysis Using xCell

To quantify immune and stromal cell type enrichment in BRCA samples and investigate immunological correlates of the four-gene panel, we applied the xCell algorithm, a gene signature-based method that deconvolutes bulk transcriptomic profiles into inferred cell-type abundances. The VST matrix (Section 2.3) was used as input, indexed by HGNC gene symbols. Expression matrices were transposed to match the expected genes x samples input format. Cell-type enrichment scores were computed using the *xCellAnalysis()* function from the xCell R package (v1.1.0) [28], yielding a matrix of normalized enrichment scores (NES) for 64 immune and stromal cell types across all samples. To assess gene-immune associations, expression profiles for the four selected genes were extracted from the VST matrix and aligned to the xCell NES matrix by sample identifier. Pairwise Spearman correlation coefficients were computed between each gene and all immune cell types using base R functions, and the resulting gene-cell-type correlation matrix was exported for visualization. The top five most correlated immune cell types (by absolute correlation) were identified per gene. Visualization was performed using Python (v3.10) and Seaborn (v0.12.2) [26], including clustered heatmaps for global patterns and lollipop plots for gene-wise top correlations. The full correlation matrix and visualizations were archived for supplementary analysis. This analysis provided insights into the immunological context of the minimal gene panel and its potential involvement in shaping or reflecting tumor immune microenvironment states.

## 3 Results

### 3.1 Transcriptome-wide differential gene expression analysis reveals widespread dysregulation in BRCA

To characterize global transcriptomic alterations associated with breast invasive carcinoma (BRCA), we performed differential gene expression analysis between tumor (n = 1,111) and normal (n = 113) samples using DESeq2. After filtering for genes with low expression and applying multiple-testing correction, we identified a total of 6,862 significantly dysregulated genes (adjusted P < 0.05, |log₂ fold change| > 1). Among these, 4,298 genes were upregulated and 2,564 genes were downregulated in tumor tissues relative to normal counterparts. The volcano plot illustrated a broad spectrum of transcriptional changes, highlighting both oncogenic activation and loss of tumor suppressive programs (Fig. 2a). A heatmap of the top 50 differentially expressed genes, ranked by adjusted P-value, further demonstrated robust expression differences that distinguish tumor from normal samples (Fig. 2b). These findings provide a comprehensive landscape of gene dysregulation in BRCA and serve as a foundation for downstream analysis for biomarker selection, and mechanistic investigation. However, the large number of differentially expressed genes underscores the need for further screening and prioritization to obtain a minimal set of biomarker genes.

**Figure 2:**
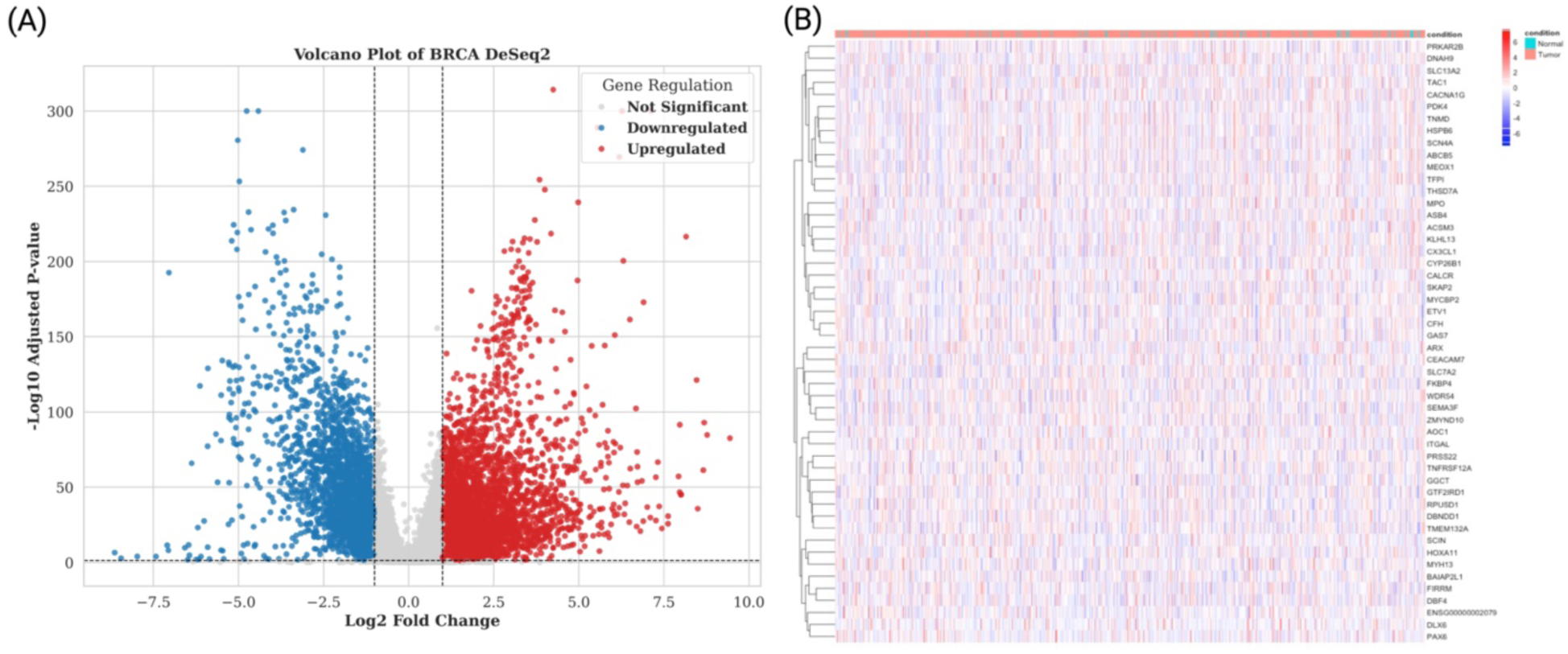
Transcriptome-wide differential expression profiling in breast cancer (BRCA). (A) Volcano plot showing differentially expressed genes between BRCA tumor (n = 1,111) and adjacent normal tissues (n = 113). Red dots represent significantly upregulated genes (adjusted P < 0.05, log₂FC > 1), while blue dots indicate significantly downregulated genes (adjusted P < 0.05, log₂FC < −1). Grey dots represent genes not meeting statistical significance thresholds. (B) Heatmap of the top 50 differentially expressed genes (ranked by adjusted P-value), based on variance-stabilized expression values. Columns correspond to individual samples, annotated by condition (tumor: red; normal: blue). Rows indicate gene symbols, and color intensity reflects expression z-scores ranging from low (blue) to high (red).

### 3.2 Global transcriptional variation in BRCA tumor and normal tissues

To explore large-scale expression heterogeneity in BRCA, we performed variance-stabilizing transformation (VST) normalization on the raw RNA-seq count matrix comprising 1,224 samples and 40,967 genes. Principal component analysis (PCA) was then applied to the normalized data to visualize global transcriptional patterns across the cohort. The resulting projection showed a clear separation between tumor (n = 1,111) and normal (n = 113) breast tissue samples along the first two principal components (PC1 and PC2), which accounted for 10.6% and 5.5% of the total variance, respectively (Fig. 3). Normal samples clustered tightly, reflecting low inter-individual variability in healthy breast transcriptomes. In contrast, tumor samples exhibited considerable spread across PCA space, indicative of extensive transcriptional reprogramming and inter-patient heterogeneity in BRCA. These findings underscore the biological distinction between malignant and normal breast tissues at the transcriptome level.

**Figure 3:**
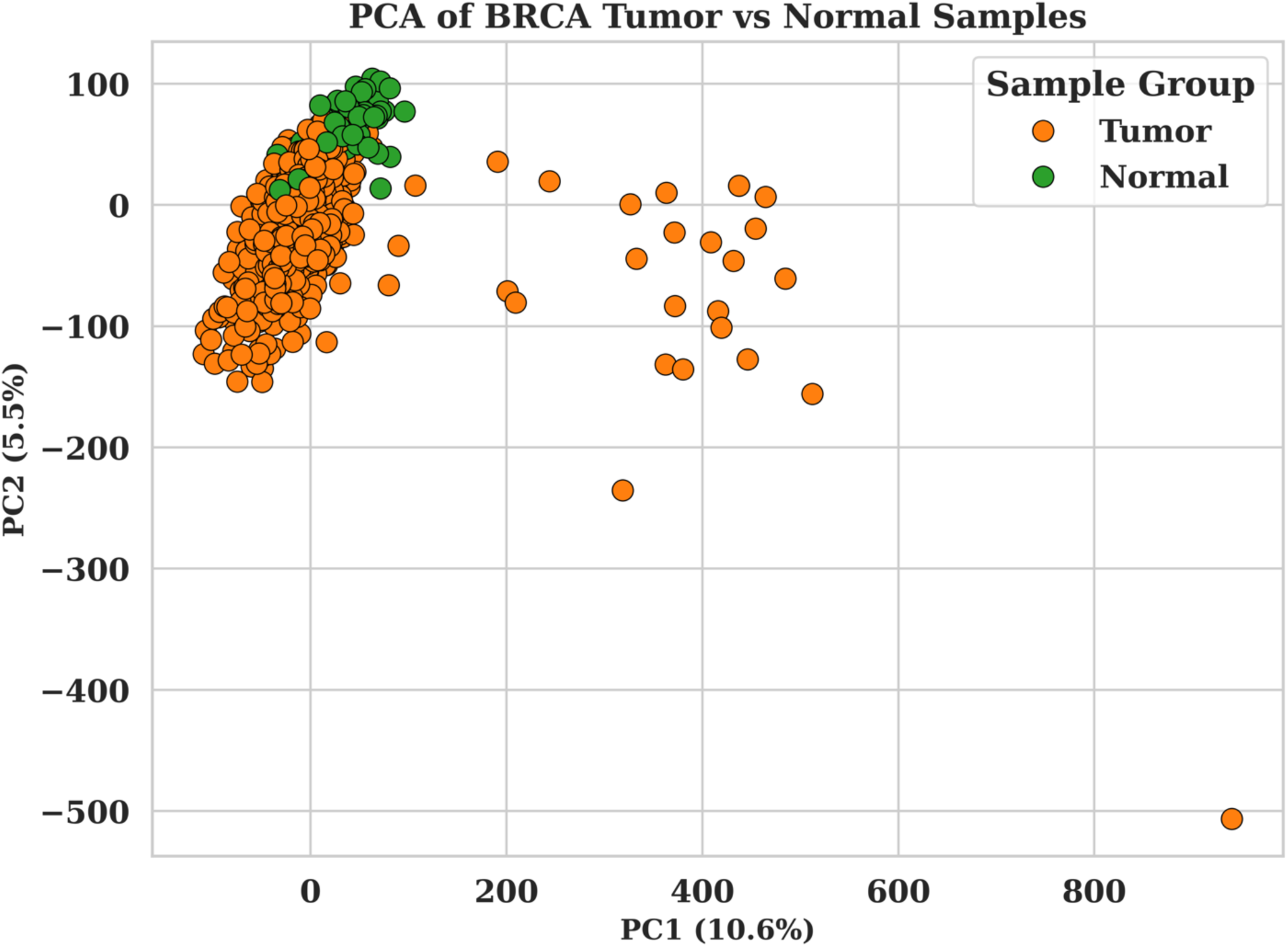
Principal component analysis (PCA) of VST-normalized RNA-seq data from BRCA tumor (orange) and normal (green) tissue samples. PC1 and PC2 explain 10.6% and 5.5% of total transcriptomic variance, respectively. Tumor samples exhibit increased dispersion, indicating greater inter-patient heterogeneity compared to normal tissues.

### 3.3 Coordinated dysregulation of ferroptosis, cuproptosis, and immune regulatory genes defines the BRCA tumor transcriptome

To investigate the convergence of cell death and immune modulation mechanisms in breast cancer (BRCA), we systematically analyzed the transcriptional landscape of ferroptosis-, cuproptosis-, and immune-associated gene sets across 1,111 tumor and 113 normal samples using variance-stabilized RNA-sequencing data. Differential expression analysis using Wilcoxon rank-sum tests revealed widespread and coordinated perturbations across all three biological axes. Among the 209 curated ferroptosis-related genes, 91 exhibited significant dysregulation (|log2FC| > 1; FDR < 0.05; Supplementary Table 2). Tumor tissues demonstrated marked upregulation of negative regulators such as EZH2 (log2FC = 2.44, FDR = 1.61×10^-40^), RRM2, and UHRF1, while several ferroptosis inducers, including GABARAPL1, CDO1, and KL, were suppressed, suggesting a concerted silencing of ferroptotic pathways in BRCA. The top dysregulated ferroptotic genes effectively separated tumor from normal samples via hierarchical clustering, with many overlapping key cell cycle and genomic instability regulators (e.g., *FOXM1, CDK1, AURKA*), reinforcing the link between ferroptosis resistance and oncogenic reprogramming. Similarly, among 133 cuproptosis-associated genes, 34 showed significant differential expression (Supplementary Table 3). Notable upregulated genes included copper transporters (*SLC25A37, COA6*), redox regulators (*LOXL1, HES6*), and transcriptional effectors such as *MYC, FOXO1*, and *FOXO4*. Established BRCA oncogenes *EGFR* and *FGF2* were also significantly dysregulated, indicating crosstalk between copper metabolism and pro-tumour growth-factor signaling pathways. The significant cuproptotic genes underscored distinct tumor-normal expression profiles. The most pronounced transcriptional reprogramming was observed in immune-related genes. Of the 21,224 unique C7 immunologic genes present in our BRCA dataset (21,373 genes in the collection), 3,699 were significantly dysregulated (|log2FC| > 1; FDR < 0.05; Supplementary Table 4), including 2,606 downregulated and 1,093 upregulated transcripts. This extensive remodeling included repression of immune effector functions and selective activation of immunomodulatory programs, indicative of immune evasion or tumor-associated immunoediting. Taken together, these findings highlight a multi-axis transcriptional reprogramming in BRCA, characterized by repression of tumor-suppressive ferroptotic and immune circuits alongside upregulation of cuproptosis- and proliferation-linked effectors. This convergence is consistent with aggressive biological behavior and therapy resistance, reflected in a ferroptosis-resistant pattern, upregulation of suppressors (*EZH2, RRM2, UHRF1*) alongside downregulation of inducers (*GABARAPL1, CDO1, KL*)[5, 8, 9], together with induction of copper transport/redox genes (*SLC25A37, COA6, LOXL1*) indicative of metabolic rewiring [10, 11, 19], dysregulation of *EGFR/FGF2* signaling linked to proliferation, angiogenesis, and immunosuppression [29–34], and broad repression of immune-effector programs (2,606 immune genes down), consistent with immune evasion [12–14, 35].

### 3.4 A distinct transcriptional footprint of ferroptosis, cuproptosis, and immune modulation in BRCA

To investigate the intersection of regulated cell death pathways and immune modulation in BRCA, we performed a comparative analysis of significantly dysregulated genes across ferroptosis-, cuproptosis-, and immune-related gene sets. Strikingly, nine genes, *EGFR, FGF2, FOXO4, IL6, CDKN2A, SLC25A37, MYC, STEAP1,* and *CP*, were found to be commonly dysregulated across all three pathways (Fig. 4; Supplementary Table 5). These genes likely represent a central axis of transcriptional crosstalk linking ferroptosis, cuproptosis, and immune signaling in the BRCA tumor microenvironment. Among these, *EGFR* (epidermal growth factor receptor) and *FGF2* (fibroblast growth factor 2) are canonical oncogenes with well-characterized roles in promoting tumor cell proliferation, angiogenesis, and immune escape [29, 30]. Their dysregulation in BRCA is consistent with sustained proliferative signaling and an immune-suppressive microenvironment [32–34]. *MYC*, a master transcriptional regulator, has been shown to repress ferroptosis and reprogram immune cell recruitment, while simultaneously enhancing metabolic reconfiguration and genomic instability hallmarks of aggressive tumors [35]. *CDKN2A*, which encodes the tumor suppressor *p16^INK4a^*, is frequently dysregulated in cancers and plays dual roles in cell cycle arrest and oxidative stress sensing [36], implicating it in both ferroptotic resistance and immune checkpoint control. Similarly, IL6 is a pleiotropic cytokine central to inflammatory signaling and immune cell polarization, whose overexpression may reflect chronic inflammation and tumor-promoting immune reprogramming [37, 38].

**Figure 4:**
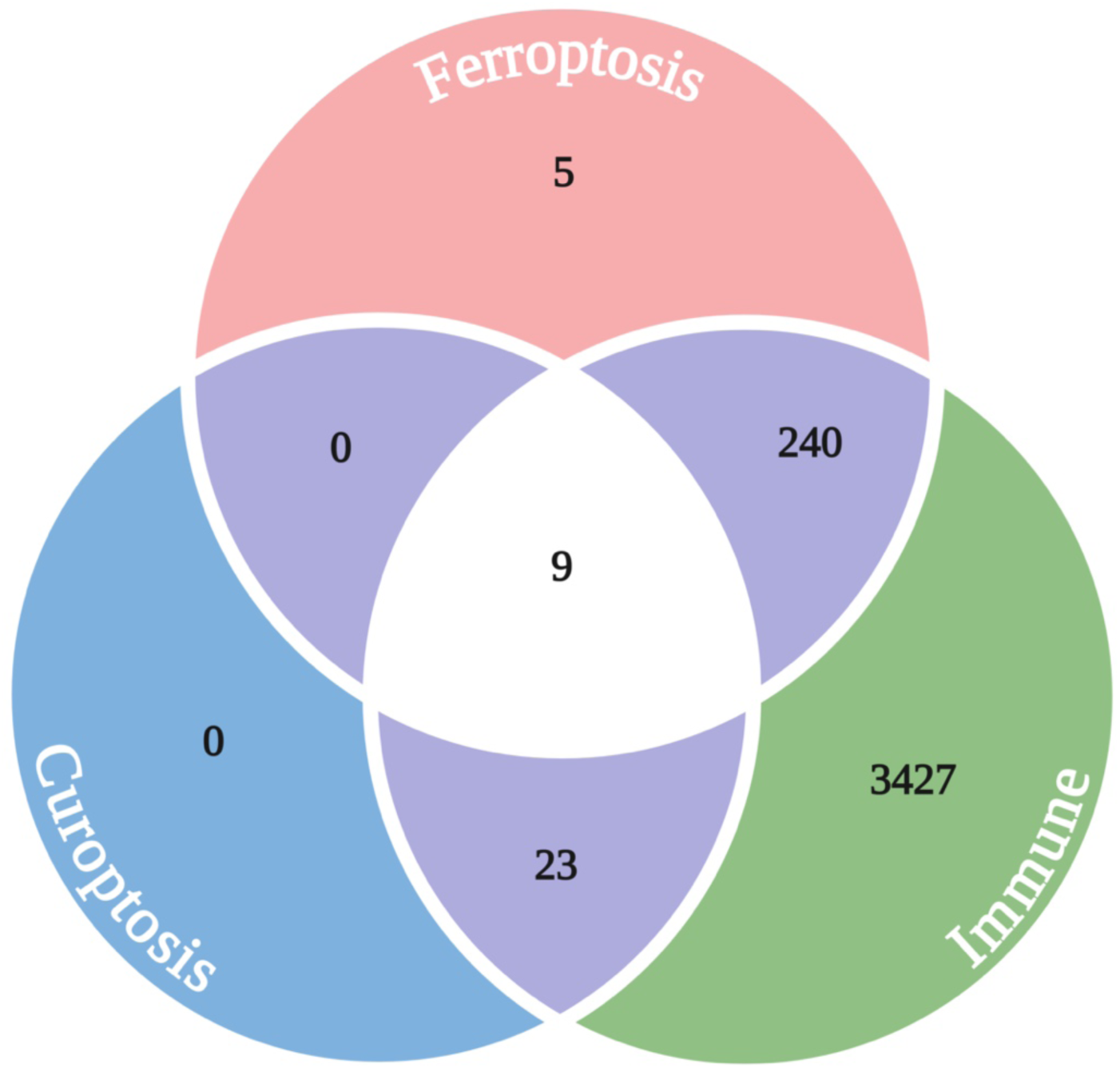
Transcriptional overlap of ferroptosis, cuproptosis, and immune-related genes in BRCA. Venn diagram showing the intersection of significantly dysregulated genes (FDR < 0.05) across ferroptosis, cuproptosis, and immune-associated gene sets in BRCA. A total of 249 genes were shared between ferroptosis and immune programs, while 32 were shared between cuproptosis and immune genes. Notably, 9 genes were shared exclusively between ferroptosis and cuproptosis, and the same nine genes were commonly dysregulated across all three pathways.

From the perspective of metal homeostasis, *SLC25A37* (also known as Mitoferrin-1) is a mitochondrial iron transporter essential for heme biosynthesis [39], while *CP* (ceruloplasmin) functions as a copper-binding ferroxidase [40], regulating both iron export and oxidative stress. Their co-dysregulation suggests a possible metabolic vulnerability wherein disrupted iron and copper handling may simultaneously affect ferroptotic and cuproptotic thresholds. STEAP1, a metalloreductase implicated in iron and copper uptake, has also been linked to immune evasion and prostate cancer aggressiveness, further supporting its role as a metal-ion-driven immune modulator [41]. *FOXO4*, a member of the FOXO transcription factor family, is known to regulate oxidative stress responses, apoptosis, and immune signaling pathways [42]. Its involvement in all three axes highlights the intricate regulation of tumor homeostasis via transcriptional feedback mechanisms that balance death and survival cues in the tumor microenvironment [43–45].

Notably, substantial overlap was observed between ferroptosis and immune-related gene expression (n = 249 genes), as well as between cuproptosis and immune pathways (n = 32 genes). Furthermore, a modest but distinct set of 9 genes was shared between ferroptosis and cuproptosis pathways independently as well as of immune involvement (Fig. 4). This observation suggests that ferroptosis, cuproptosis, and immune programs exhibit transcriptional integration in BRCA, challenging the notion of its functioning as a fully discrete axis and instead indicating a more nuanced landscape of pathway crosstalk.

Together, these nine-gene signature defines a biologically plausible regulatory core that integrates oxidative stress responses, metal ion metabolism, and immune signaling in BRCA. Furthermore, this intersection of multiple pathways known for their dysregulation in breast cancer can be leveraged to generate a diagnostic panel of the disease, as it is highly likely that central genes will amplify transcriptional signal resulting from the pathological state.

### 3.5 Machine learning prioritization reveals a compact four-gene panel for BRCA stratification

To derive a diagnostically efficient and mechanistically grounded subset from the nine-gene intersection panel (*EGFR, FGF2, FOXO4, IL6, CDKN2A, SLC25A37, MYC, STEAP1, CP*), we implemented a multi-model machine learning framework incorporating both statistical regularization and interpretable feature attribution. Expression data for these genes were extracted from the VST-normalized matrix and z-transformed. Using binary tumor vs. normal labels from TCGA metadata, we trained LASSO logistic regression, Random Forest (RF), and XGBoost classifiers on the 9-gene panel. Gene prioritization was performed via L1-penalized coefficients (LASSO), impurity-based feature importance (RF), and gain scores (XGBoost), followed by global SHAP (SHapley Additive exPlanations) value analysis on the XGBoost model to quantify model-agnostic contributions (Fig. 5A-C; Supplementary Table 6). Concordantly across models, four genes, *FOXO4, EGFR, FGF2*, and *CDKN2A*, emerged as the top-ranked features (Fig. 5, Supplementary Table 6). *FOXO4*, a master regulator of oxidative stress and apoptosis, showed the strongest negative LASSO coefficient and the highest SHAP impact. *EGFR* and *FGF2* are canonical oncogenes that regulate proliferation and immune evasion, while *CDKN2A* modulates both cell cycle checkpoints and redox-dependent stress signaling, consistent with dual roles in ferroptotic and immune pathways. SHAP beeswarm and waterfall plots further demonstrated the consistent directionality and individualized impact of these features across BRCA samples (Fig. 5B-C). Notably, this core quartet captured over 97% of the maximal predictive accuracy across models (Fig. 5E; Supplementary Table 7), indicating that this reduced gene set retained both biological relevance and diagnostic fidelity. The importance of dimensionality reduction extends beyond interpretability and visualization; it enables practical translation into clinical assays by minimizing redundancy and maximizing biomarker efficiency. Furthermore, by aligning feature prioritization with functional relevance across ferroptosis, cuproptosis, and immune regulation, this four-gene panel offers a compact molecular signature with potential utility in biomarker development and therapeutic targeting. To further validate the discriminative power of the selected four-gene panel, we performed principal component analysis (PCA) using z-transformed VST expression values across BRCA tumor and normal samples. The PCA projection demonstrated clear group separation, with PC1 and PC2 jointly explaining ∼70% of total variance (Fig. 5F). In parallel, violin plots showed robust and consistent differential expression between tumor and normal samples for each of the four genes (Fig. 5G), highlighting their potential as clinically relevant biomarkers.

**Figure 5:**
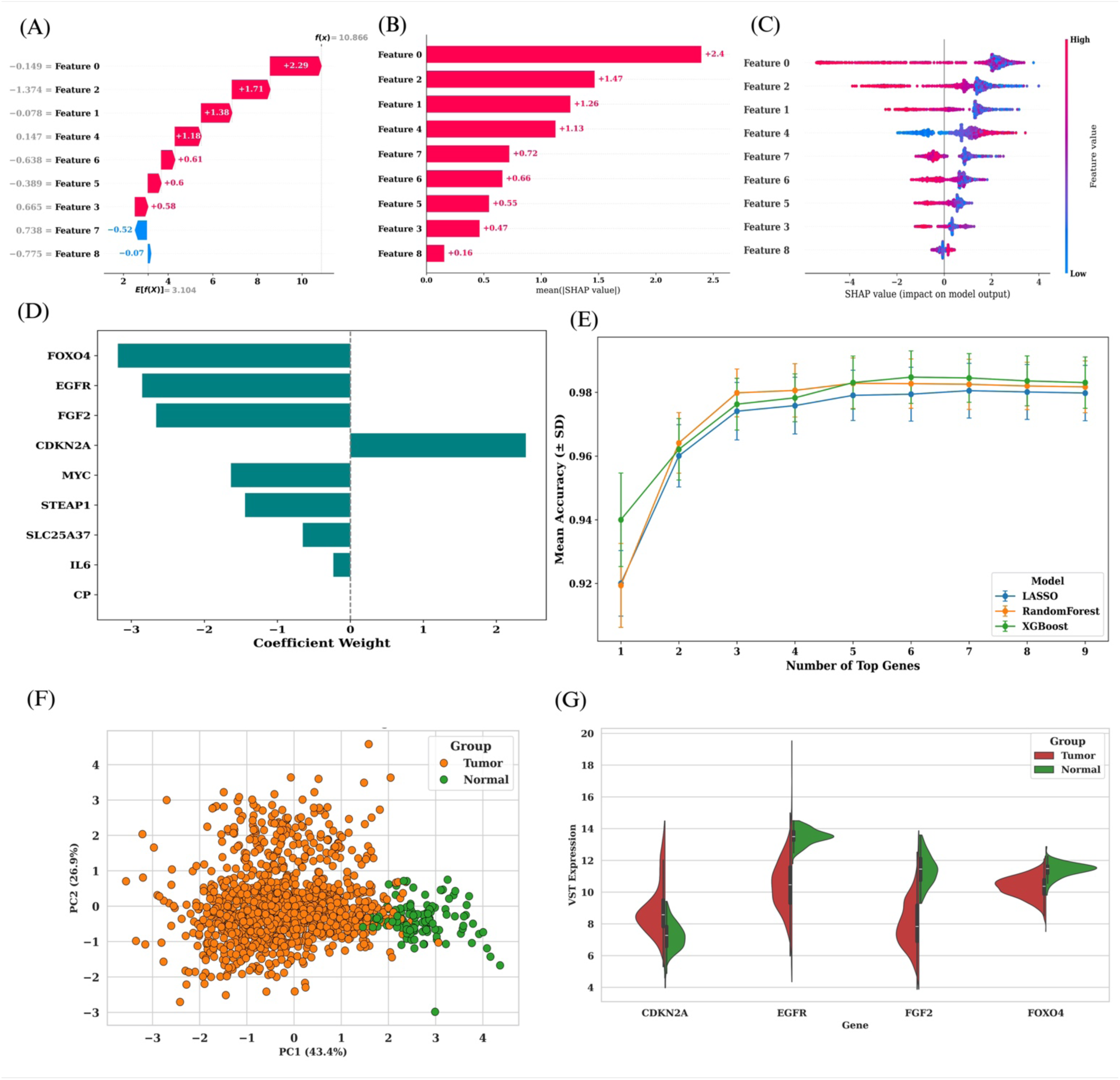
Prioritization and dimensionality reduction of a 9-gene panel into a core 4-gene BRCA classifier. **(A)** SHAP (SHapley Additive exPlanations) waterfall plot illustrating the contribution of each feature to an individual BRCA sample’s XGBoost model prediction. Feature 0 (*FOXO4*) and Feature 2 (*EGFR*) exhibit the highest positive impacts on the model output. (B) Mean absolute SHAP values across all samples ranked by importance, highlighting the dominant contribution of *FOXO4, EGFR, FGF2,* and *CDKN2A*. (C) SHAP beeswarm plot showing the distribution and directionality of SHAP values across the cohort. Red indicates high expression and blue low expression, reinforcing *FOXO4, EGFR, FGF2,* and *CDKN2A* as top features with consistent predictive direction. (D) LASSO logistic regression coefficient weights for the 9-gene panel. *FOXO4* shows the strongest negative weight, supporting its role as a critical tumor suppressive classifier. (E) Bootstrap-averaged mean accuracy curves for LASSO, Random Forest, and XGBoost models trained on increasing numbers of top-ranked genes. The core four genes (*FOXO4, EGFR, FGF2, CDKN2A*) capture >97% of peak classification performance, supporting their utility as a minimal, mechanistically informed biomarker panel. (F) PCA of BRCA tumor vs. normal samples based on z-transformed VST expression of *FOXO4, EGFR, FGF2*, and *CDKN2A*. Tumor and normal samples show clear separation, with PC1 and PC2 accounting for 43.4% and 26.9% of variance, respectively. (G) Violin plots comparing gene expression levels across tumor and normal tissues. All four genes exhibit consistent differential expression, reinforcing their relevance to BRCA biology and classification.

To evaluate the ability of our four-gene panel (*FOXO4, EGFR, FGF2,* and *CDKN2A*) to discriminate between BRCA tumors and adjacent normal tissues, we performed dimensionality reduction and visualization of expression data. When projected into two dimensions using principal component analysis (PCA), the full transcriptome showed partial overlap between tumor and normal samples, reflecting the high heterogeneity of BRCA (Fig. 5F). In contrast, the reduced model based only on the four selected genes yielded markedly improved separation, with tumors clustering distinctly away from normal tissues (Fig. 6). This result demonstrates that the four-gene panel captures the most informative expression shifts underlying tumorigenesis, providing a biologically grounded yet compact classifier that enhances interpretability while maintaining strong discriminatory power.

**Figure 6:**
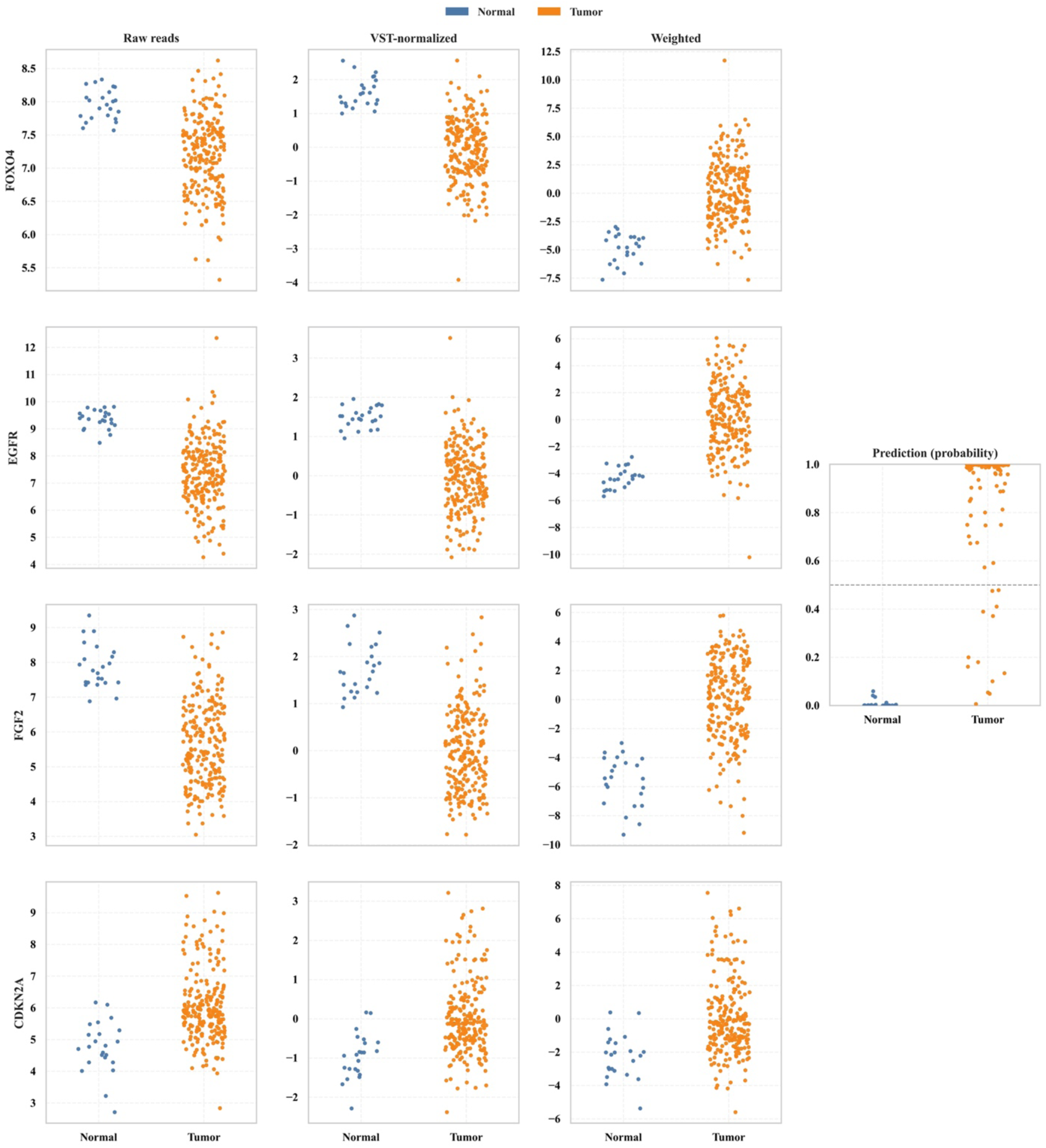
Improved separation of BRCA tumor and normal samples by the four-gene panel using the test set. The plots show the compact model achieves clearer discrimination between tumor (orange) and normal (blue) samples, highlighting its ability to condense complex transcriptional alterations into a mechanistically interpretable signature.

### 3.6 Robust diagnostic classification of BRCA using a minimal four-gene panel across diverse machine learning algorithms

To evaluate the generalizability and clinical potential of the minimal four-gene panel *(FOXO4, EGFR, FGF2, CDKN2A)*, we benchmarked its predictive performance across a suite of 11 commonly employed classification algorithms using a bootstrapped cross-validation framework (n = 100 iterations). Each model was trained using a SMOTE-augmented pipeline to mitigate class imbalance and assessed on both training and test sets via multiple evaluation metrics (Supplementary Table 8). Overall, all models demonstrated high classification fidelity on the test data, with mean accuracies exceeding 96% across the board (Fig. 7). Ensemble methods consistently performed at the upper bound of accuracy: XGBoost (97.9% ± 0.008), Random Forest (97.8% ± 0.009), and AdaBoost (97.6% ± 0.009). These models also achieved high F1 scores (≥ 0.986) and area under the ROC curve (AUC > 0.99), underscoring their ability to simultaneously capture sensitivity and precision. Notably, XGBoost achieved the highest test AUC (0.995 ± 0.0038), further corroborating its utility in high-confidence diagnostic prediction. Linear models, including LASSO (96.6% ± 0.011) and Logistic Regression (96.6% ± 0.011), offered slightly lower but still robust performance while maintaining model interpretability. Support Vector Machine (SVM) and Artificial Neural Networks (ANN) also achieved strong predictive metrics (accuracy > 97.1%; F1 score > 0.983), indicating the versatility of the panel across both linear and non-linear classifiers. Interestingly, even relatively simpler models such as Naive Bayes and Decision Tree achieved classification accuracies of 97.4% and 97.1%, respectively, highlighting the discriminative strength of this four-gene signature. Importantly, ROC AUC, sensitivity, and specificity values remained consistently high across classifiers, indicating that performance was not skewed toward either tumor or normal samples. The relatively low standard deviations across bootstraps (e.g., accuracy std < 0.01) further confirmed the robustness and reproducibility of predictions (Supplementary Table 7). These results collectively demonstrate that the *FOXO4, EGFR, FGF2,* and *CDKN2A* panel provides a compact yet highly discriminative molecular signature capable of classifying BRCA tumors with high accuracy and reproducibility across diverse modeling strategies. The minimal size of the panel, combined with its strong diagnostic performance, supports its translational utility in clinical molecular diagnostics and resource-constrained screening pipelines.

**Figure 7:**
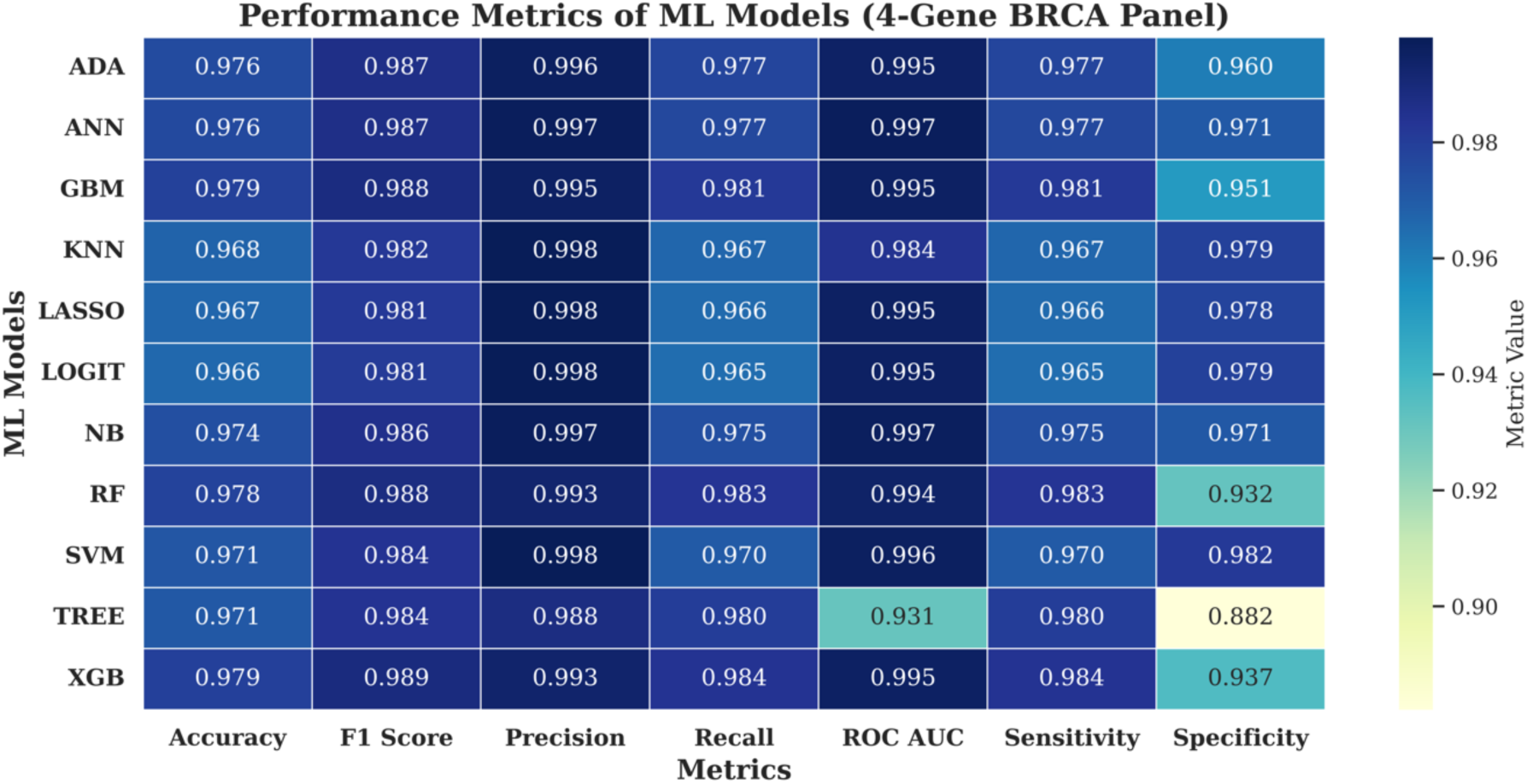
Performance evaluation of machine learning models trained on the minimal 4-gene BRCA panel. Heatmap summarizing the average classification performance of 11 machine learning models trained using *FOXO4, EGFR, FGF2,* and *CDKN2A* expression values as features. Metrics were computed on test sets across 100 bootstrapped splits and include Accuracy, F1 Score, Precision, Recall, ROC AUC, Sensitivity, and Specificity. Darker shades indicate higher metric values, with uniformly strong performance observed across all classifiers. XGBoost (XGB), Random Forest (RF), Gradient Boosting (GBM), and AdaBoost (ADA) showed consistently high accuracy (≥0.976), while SVM and LASSO achieved superior sensitivity and specificity balance. These results validate the predictive robustness of the compact 4-gene panel across diverse classification paradigms.

### 3.7 Immune correlates of the four-gene panel highlight distinct tumor microenvironment interactions

To characterize the immunological landscape associated with the four-gene panel, we employed the xCell algorithm to estimate enrichment scores for 64 immune and stromal cell types across 1,224 BRCA samples using VST-normalized expression profiles. xCell deconvolution yielded normalized enrichment scores (NES), which were subsequently correlated with gene expression levels of *FGF2, EGFR, FOXO4,* and *CDKN2A* using Spearman’s method. This analysis revealed distinct, gene-specific immune microenvironment associations (Fig. 8; Supplementary Table 11). FGF2 displayed strong positive correlations with stromal components, including megakaryocytes (ρ = 0.62), StromaScore (ρ = 0.59), HSCs (ρ = 0.56), and fibroblasts (ρ = 0.53), while showing moderate negative associations with Th1 cells (ρ = −0.55), suggesting a link with tumor fibrosis and reduced cytotoxic immunity. *EGFR* correlated positively with epithelial-like signatures, including keratinocytes (ρ = 0.63) and astrocytes (ρ = 0.52), while being negatively associated with osteoblasts and common lymphoid progenitors, highlighting its potential role in epithelial remodeling rather than immune evasion. *FOXO4* showed modest correlations with both macrophages (ρ = 0.31) and Th2 cells (ρ = −0.40), consistent with its known dual roles in immune tolerance and oxidative stress. Among the four genes, *CDKN2A* showed the broadest and most consistent immune associations, with moderate positive correlations across multiple lineages, e.g., plasma cells (ρ = 0.39), M1 macrophages (ρ = 0.33), and aDCs (ρ = 0.29), supporting a role in shaping both innate and adaptive immune responses. All top five correlations per gene were statistically significant (*P* < 0.01, FDR-adjusted), indicating robust associations unlikely to occur by chance. The lollipop plots (Fig. 8) illustrate these gene-cell-type relationships, revealing a convergent enrichment in stromal and immune cell types for most genes, particularly *FGF2* and *CDKN2A*. Collectively, these findings suggest that the four-gene panel is not only predictive but also immunologically anchored, capable of capturing variations in the tumor immune microenvironment. This further supports its utility as a stratification biomarker in breast cancer and a candidate for immunogenomic hypothesis generation.

**Figure 8:**
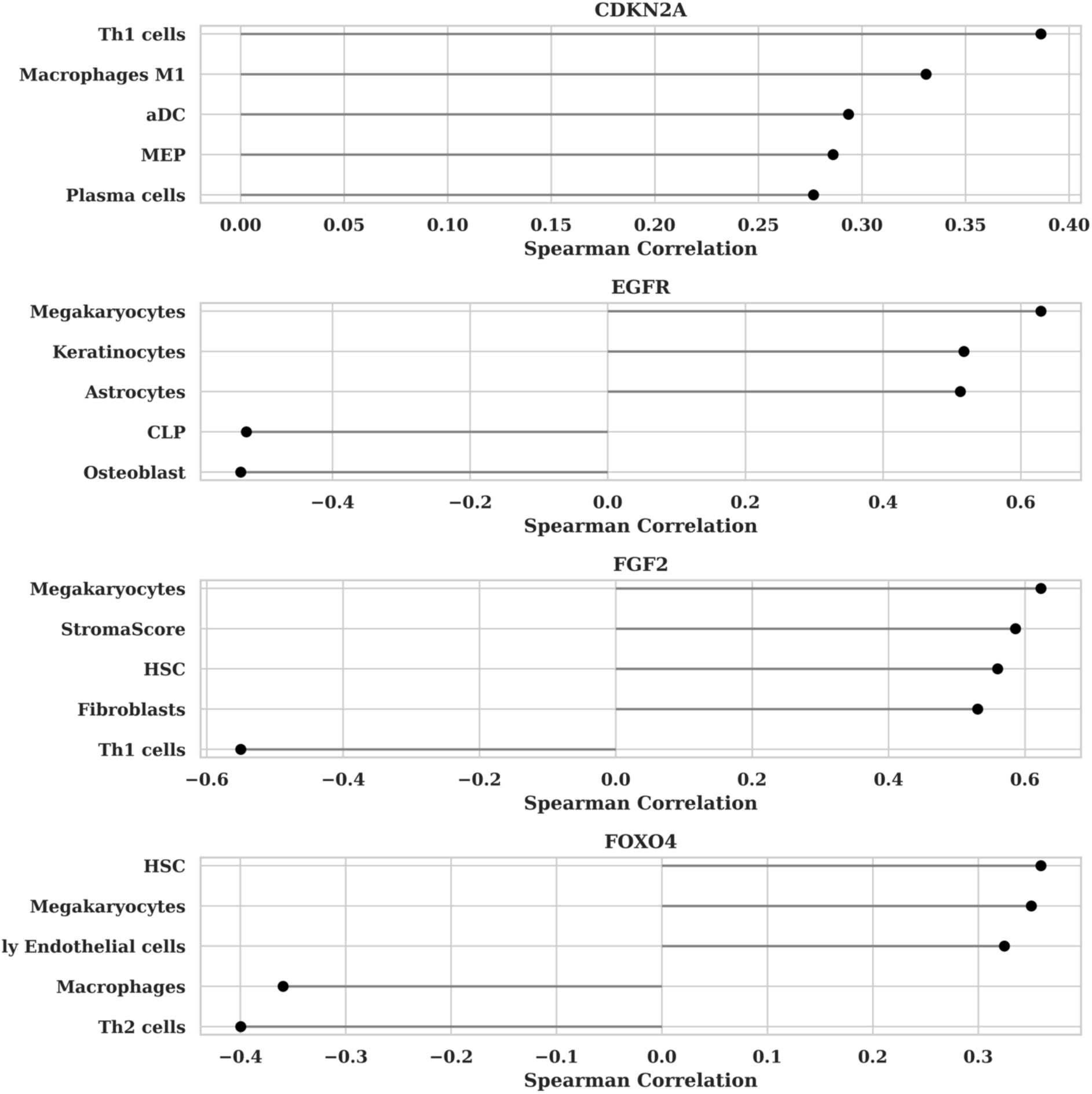
Immune cell-type correlations for the four-gene panel in BRCA. Lollipop plots showing the top five immune or stromal cell types (xCell-inferred) most strongly correlated with the expression of each gene in the four-gene panel (*CDKN2A, EGFR, FGF2, FOXO4*). Spearman correlation coefficients were computed across BRCA samples between gene expression and cell-type enrichment scores. Positive and negative associations are visualized along the x-axis. These results highlight distinct immunological contexts for each gene, with *FGF2* and *CDKN2A* displaying strong associations with stromal and innate immune cell populations.

## 4 Discussion

Despite decades of progress in molecular profiling and personalized therapy, breast cancer (BRCA) remains clinically challenging due to its pronounced molecular heterogeneity and complex immune-metabolic interactions. Here, we present a biologically anchored, machine learning-prioritized four-gene panel, *FOXO4, EGFR, FGF2,* and *CDKN2A*, that integrates ferroptosis, cuproptosis, and immune signaling axes into a single compact panel. Unlike traditional transcriptomic signatures that prioritize statistical discriminability over functional relevance, our panel was derived from convergent dysregulation across regulated cell death and immune programs and refined through multi-model interpretability metrics. This approach yielded a panel of genes that demonstrates high predictive performance across diverse machine learning models. Importantly, our results indicate that no single gene in this panel is sufficient to accurately predict BRCA status on its own. Each gene contributes only partial discriminative power, reflecting distinct biological axes such as proliferation (*EGFR, FGF2*), cell cycle control (*CDKN2A*), and stress signaling (*FOXO4*). It is only through their combined weights in the model that robust separation between tumor and normal samples is achieved, underscoring the necessity of a multi-gene integration strategy for reliable classification. Table 1 provides an overview of the biological functions, regulatory context, and involvement in ferroptosis, cuproptosis, and immune pathways for each gene in the panel.

**Table 1:** Functional and clinical characterization of the four-gene BRCA panel. Overview of expression trends, BRCA-specific roles, relevance in other cancers, regulatory mechanisms, therapeutic targeting status, and involvement in ferroptosis, cuproptosis, and immune pathways for FOXO4, EGFR, FGF2, and CDKN2A.

| Gene | *FOXO4* | *EGFR* | *FGF2* | *CDKN2A* |
| --- | --- | --- | --- | --- |
| Expression in this study | Downregulated | Downregulated | Downregulated | Upregulated |
| Known Function in BRCA | Tumor suppressor; regulates apoptosis and oxidative stress via PI3K/AKT inhibition [46, 47]. | Canonical oncogene; overexpressed in ~30% of BRCA (especially TNBC); drives proliferation [48, 49] | Pro-angiogenic factor; involved in BRCA invasion and endocrine resistance [50]. | Tumor suppressor; controls G1/S checkpoint via p16INK4a and p14ARF; inactivated in many BRCA subtypes [51]. |
| Roles in Other Cancers | Tumor suppressor in colon [45], prostate [52], and glioblastoma [44]. | Targeted in NSCLC [53], glioblastoma [54], and colorectal cancer [55]. | Oncogenic in prostate [56, 57], lung [58, 59], and gastric cancers [60]. | Deleted or methylated in melanoma [61, 62], pancreatic [63, 64], and bladder cancers [65, 66]. |
| **Regulation** | Regulated by AKT, SIRT1, and miRNAs (e.g., miR-27a) [67-69]. | Transcriptionally regulated by HIF1α, AP-2, and miRNAs (e.g., miR-145) [70-72]. | Induced by hypoxia [73], HIF1α [74], and post-transcriptional regulation via miR-497 [75]. | Transcriptionally repressed by EZH2 [76, 77]; activated via DNA damage; miR-24 [78], miR-106b regulate it [79]. |
| **Clinical Targeting / Approved Drugs** | No FDA-approved drugs, but being explored in pro-apoptotic peptide-based therapies (e.g., FOXO4-DRI) [80]. | FDA-approved inhibitors: erlotinib [81], gefitinib [82, 83], osimertinib [84, 85]. | Targeted in preclinical trials (e.g., FGFR inhibitors) [86]. | No direct drugs, but reactivation strategies via EZH2 inhibitors under study [87, 88]. |
| **Involvement in Ferroptosis / Cuproptosis / Immune Pathways** | **Ferroptosis:** Interacts with SLC7A1 regulation [43]; **Immune:** Modulates interferon responses via ROS [89], PD-L1 and VEGF [90]; **Cuproptosis:** No direct mechanistic evidence; potential unexplored involvement based on co-expression in BRCA. | **Immune:** EGFR suppresses anti-tumor immunity via PD-L1 upregulation [31]; **Ferroptosis:** EGFR inhibition sensitizes to ferroptosis via GPX4 depletion [91, 92]; **Cuproptosis:** Emerging link with copper metabolism [93]. | **Immune:** Promotes M2 macrophage polarization and immune escape [94]; **Ferroptosis:** Linked to NRF2 signaling and iron homeostasis [95, 96]; **Cuproptosis:** Emerging role in copper metabolism [97]. | **Ferroptosis:** Regulates sensitivity independently of p53; modulates lipid peroxidation susceptibility [98, 99]; **Immune:** Correlated with immune checkpoint expression in some tumors [100, 101]; **Cuproptosis:** emerging evidences suggest its role in copper metabolism pathways in different cancers [102-104] |

Each gene within the refined panel contributes uniquely to the oncogenic and immunological landscape of BRCA, reinforcing the panel’s functional coherence. *FOXO4*, a transcription factor best known for orchestrating oxidative stress responses and cell fate decisions, emerged as a central node in our framework [46, 47]. Its consistent downregulation in BRCA samples, coupled with moderate correlations to interferon signaling and immune cell infiltration, suggests an underexplored role in modulating immune surveillance and cell death susceptibility, potentially via redox imbalance and ferroptosis resistance [43, 89, 90]. *EGFR*, a canonical oncogene and established therapeutic target, exhibited widespread dysregulation across BRCA subtypes and was strongly associated with epithelial remodeling and immune checkpoint activation [31, 48, 49, 53, 91–93]. Its inclusion underscores the convergence of growth factor signaling with ferroptotic suppression and immune escape. *FGF2*, another mitogenic driver, revealed pro-angiogenic and pro-stromal transcriptional signatures, aligning with its known function in vascular remodeling and tumor fibrosis, while simultaneously displaying ferroptosis-related regulatory features through NRF2 signaling and iron metabolism [50, 95–97]. Finally, *CDKN2A*, frequently dysregulated in BRCA and multiple epithelial cancers, plays dual roles in G1/S checkpoint control and oxidative stress regulation [51]. Its upregulation in this context may reflect a compensatory response to cell cycle dysregulation or immune-editing pressure [98–104]. Collectively, these genes offer more than classification; they represent biologically interpretable readouts of the tumor’s redox state, proliferative potential, and immune status as shown in Fig 9.

**Figure 9:**
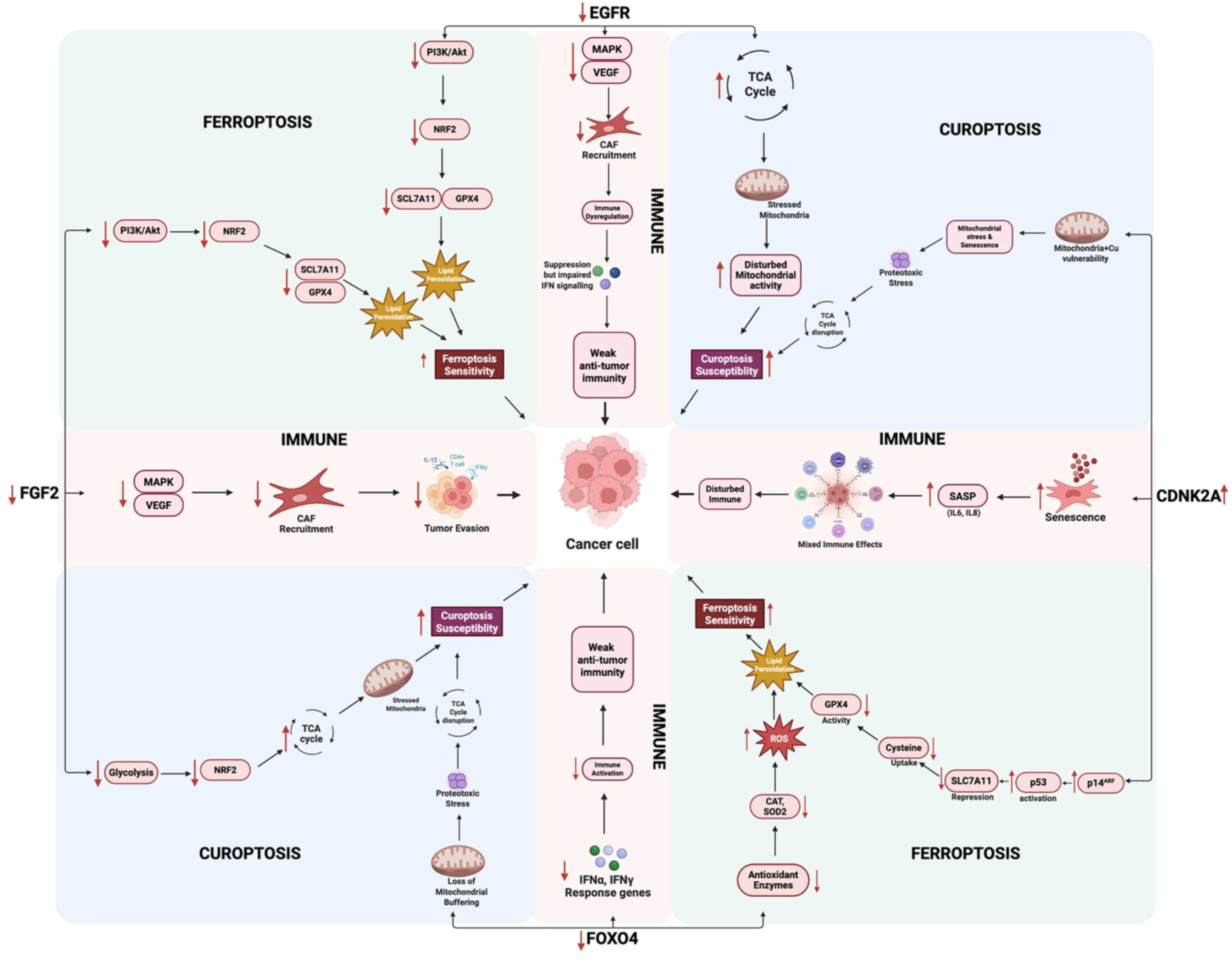
Integrated pathway map of EGFR, FGF2, FOXO4, and CDKN2A dysregulation in BRCA. Downregulation of EGFR, FGF2, FOXO4, and upregulation of CDKN2A converge on three major pathways in breast cancer: ferroptosis (via loss of antioxidant defenses and SLC7A11 repression), cuproptosis (via increased TCA cycle reliance and mitochondrial stress), and immune modulation (via reduced VEGF/CAF signaling, impaired interferon responses, and SASP-driven inflammation). Together, these events reprogram BRCA cells toward altered death thresholds and immune evasion.

A key innovation of this study lies in the conceptual convergence of ferroptosis, cuproptosis, and immune signaling, three mechanistically distinct yet transcriptionally intersecting pathways that shape tumor progression and therapeutic response [105]. Historically treated as independent axes, our integrative analysis revealed a coordinated dysregulation of these programs in BRCA, with nine genes, including *FOXO4, EGFR, FGF2,* and *CDKN2A*, shared across all three regulatory domains. This overlap was not merely incidental; it corresponded to functionally meaningful roles in redox regulation, metal ion metabolism, and immune interface. *FOXO4* and *CDKN2A* exhibited cross-talk between oxidative stress sensing and immune checkpoint pathways, while EGFR and *FGF2* bridged growth factor signaling with ferroptotic resistance and immunosuppressive remodeling. Although cuproptosis remains a nascent concept in cancer biology, our findings suggest that copper-related transcriptional reprogramming intersects with canonical oncogenic circuits, warranting further exploration. Importantly, the co-dysregulation of these pathways suggests that BRCA tumors do not simply silence or activate individual death mechanisms, but rather rewire a broader regulatory network that balances immune evasion, metabolic adaptation, and cell death thresholds.

Beyond its mechanistic grounding, the four-gene panel demonstrated robust and reproducible diagnostic performance across a diverse suite of machine learning classifiers, achieving consistently high accuracy, precision, and AUC values. Ensemble-based models such as XGBoost and Random Forest reached test accuracies exceeding 97%, while even simpler algorithms such as logistic regression and decision trees yielded comparable performance, highlighting the panel’s adaptability across computational platforms. Consistent with this, single-gene models underperformed substantially compared to the four-gene combination, underscoring that predictive fidelity arises from their complementary weights rather than any single dominant contributor. Dimensionality reduction from the initial nine-gene intersection to a compact four-gene core panel guided not only by statistical feature importance but also by SHAP-based interpretability, ensuring that the retained genes preserved high predictive performance while remaining biologically grounded in ferroptosis, cuproptosis, and immune pathways. This approach enhances the panel’s clinical feasibility, reducing cost, complexity, and technical barriers associated with larger multigene assays, particularly in resource-limited or point-of-care settings. Moreover, the use of only four genes facilitates downstream validation via qRT-PCR, IHC, or targeted panels, streamlining potential integration into existing diagnostic workflows. Taken together, the panel’s diagnostic fidelity, interpretability, and practicality position it as a compelling candidate for translational deployment in BRCA stratification and surveillance.

While our findings provide compelling evidence for the diagnostic and mechanistic value of the four-gene panel, several limitations merit consideration. While cuproptosis is a relatively nascent pathway with evolving definitions, our integration of this axis into BRCA stratification represents a conceptual advance that invites experimental investigation. Future work should aim to validate the panel across independent clinical cohorts, assess its prognostic utility, and explore its role in predicting response to ferroptosis- and copper-targeting agents. This study establishes a biologically interpretable, computationally robust, and clinically feasible four-gene classifier that integrates ferroptosis, cuproptosis, and immune regulation into a unified diagnostic framework. By grounding molecular classification in core oncogenic and cell death processes, our approach advances precision oncology efforts toward minimal, mechanism-informed, and therapeutically actionable biomarkers in breast cancer.

## 5 Conclusion

This study introduces a minimal yet mechanistically grounded four-gene panel, *FOXO4, EGFR, FGF2,* and *CDKN2A*, that captures convergent dysregulation across ferroptosis, cuproptosis, and immune signaling in breast cancer. Through integrative transcriptomic analysis, machine learning-based prioritization, and multi-modal validation, we demonstrate that this panel not only enables high-fidelity tumor identification but also reflects core oncogenic and immunological processes. Its compact design supports clinical feasibility, while its functional relevance highlights therapeutic opportunities across established and emerging pathways. By bridging cell death regulation with tumor immune contexture, our framework offers a translationally actionable strategy for biologically informed BRCA stratification.

## List of Abbreviations

AUC: Area under the receiver operating characteristic curve
BRCA: Breast cancer
CDKN2A: Cyclin dependent kinase inhibitor 2A
CV: Cross-validation
EGFR: Epidermal growth factor receptor
FGF2: Fibroblast growth factor 2
FOXO4: Forkhead box O4
LASSO: Least absolute shrinkage and selection operator regression (logistic regression with L1 penalty)
ML: Machine learning
RF: Random Forest classifier
ROC: Receiver operating characteristic
SHAP: Shapley additive explanations
SMOTE: Synthetic minority oversampling technique
SVM: Support vector machine
TCGA: The Cancer Genome Atlas
XGBoost: Extreme gradient boosting classifier

## Declarations

### Ethics approval and consent to participate

Not applicable. This study used only publicly available datasets (TCGA-BRCA) and did not involve human participants, human tissue, or animal experiments.

### Consent for publication

Not applicable. This manuscript does not include data from any individual person.

### Availability of data and materials

The RNA-sequencing and clinical data for breast cancer (BRCA) analyzed in this study were obtained from The Cancer Genome Atlas (TCGA) via the Genomic Data Commons (GDC) portal (https://portal.gdc.cancer.gov/). All processed datasets generated during this study are provided in the Supplementary Data. Custom R and Python scripts used for data processing, statistical analysis, and model development are available from the corresponding author on reasonable request.

### Authors’ contributions

- **S.A.S. (Syed Ahsan Shahid):** Conceptualization, data analysis, machine learning model development, manuscript drafting.
- **A.A.H. (Professor Ahmed Al-Harrasi):** Supervision, critical revision of the manuscript.
- **A.A.S. (Dr. Adil Al-Siyabi):** Supervision, project administration, manuscript editing. All authors read and approved the final manuscript.

## Acknowledgements

The authors gratefully acknowledge the University of Nizwa, Natural and Medical Sciences Research Center, for providing computational and infrastructural support.

